## Supplemental Figures for "Efficacy of epetraborole against *Mycobacterium abscessus* is increased with norvaline"

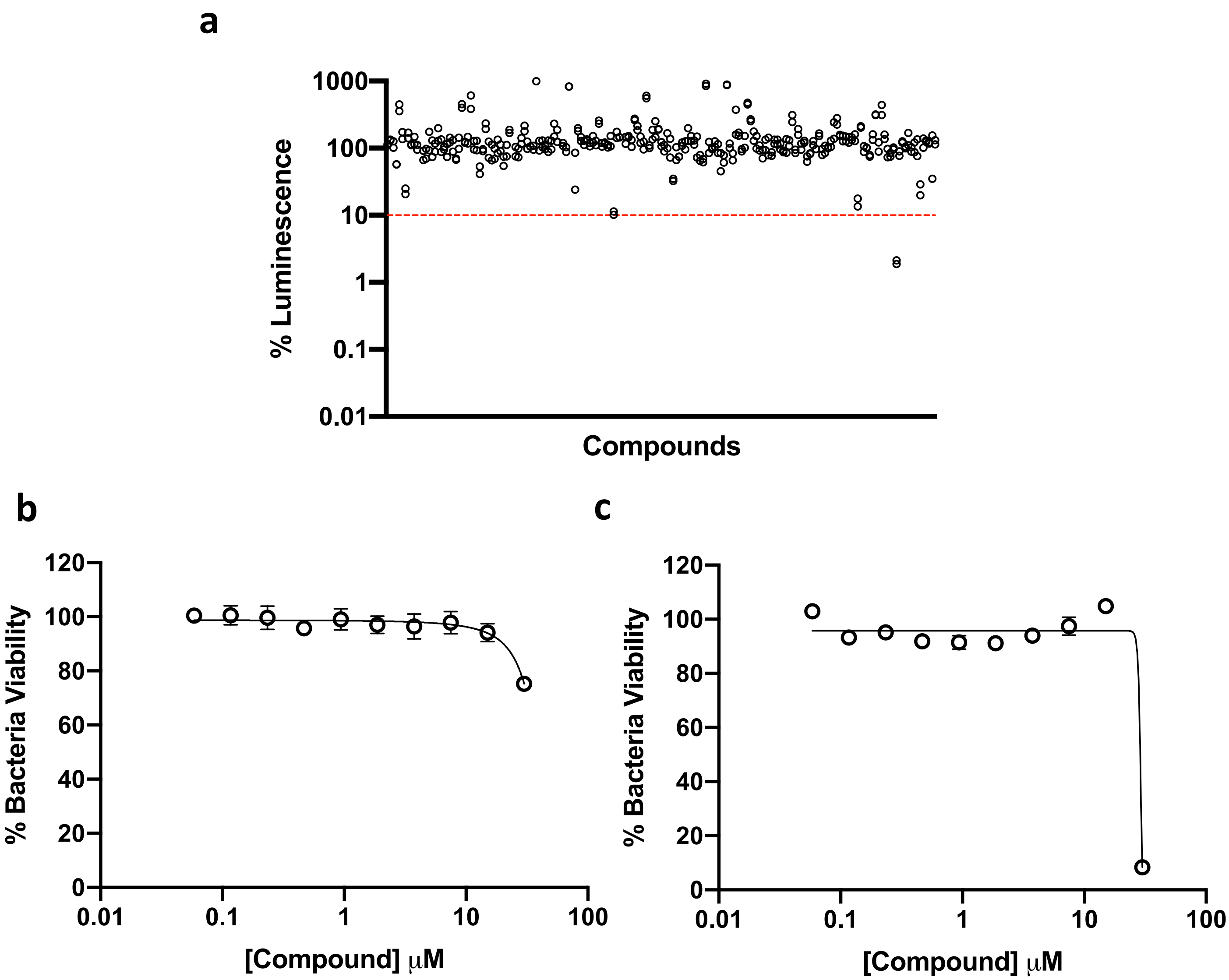


**Supplementary Figure 1.**
*M. abscessus* phenotypic screening with 176 TB-active compounds from GSK. **a** Luminescence relative to drug free conditions using *M. abscessus* lux. Data is shown in duplicate. **b-c** Secondary screening of primary hits using REMA on *M. abscessus* ATCC 19977. % bacteria viability relative to drug free conditions. Data is shown as mean ± SD from technical triplicates. Dashed red line indicates 10% viability threshold when screened at 10μM. Three hits were identified from primary screen however, all three did not pass the secondary screen. No active compounds were identified from this library.

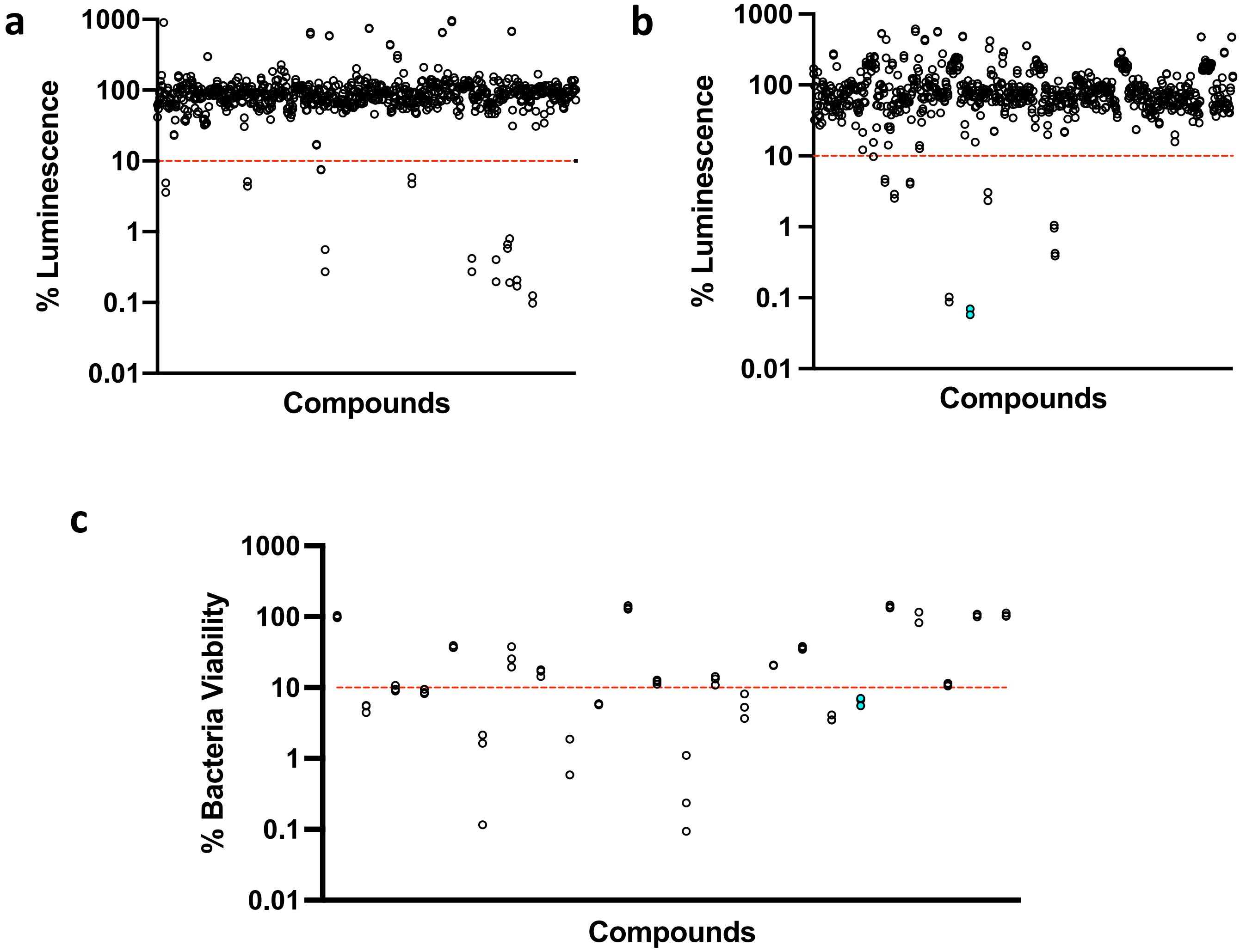


**Supplementary Figure 2.**
*M. abscessus* ATCC 19977 phenotypic screening of compounds from MMV. **a** Pathogen box; luminescence relative to drug free conditions using *M. abscessus* lux. Data is shown in duplicate. **b** Pandemic response box; luminescence relative to drug free conditions using *M. abscessus* lux. Data is shown in duplicate. **c** Secondary screening of primary hits using REMA on *M. abscessus* ATCC 19977. % bacteria viability relative to drug free conditions. Data is shown in technical triplicate. Dashed red line indicates 10% viability threshold when screened at 10$\mu$M. 20 compounds passed the primary screen and 14 passed the secondary screen. Only three compounds were still active after acquiring fresh batch and displayed dose-dependent activity (3/800). EPT in cyan.

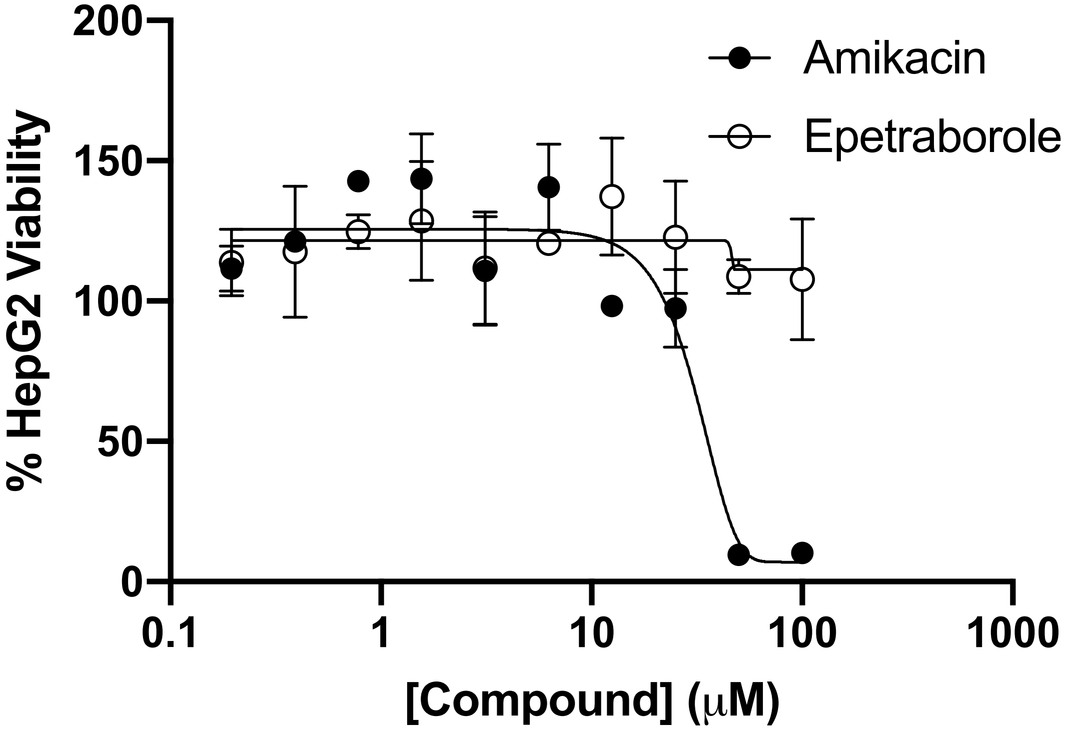


**Supplementary Figure 3.**
Cytotoxicity of EPT (white circles) and AMK (black circles) on HepG2 cells for three days. Data is mean ± SD from technical triplicates.

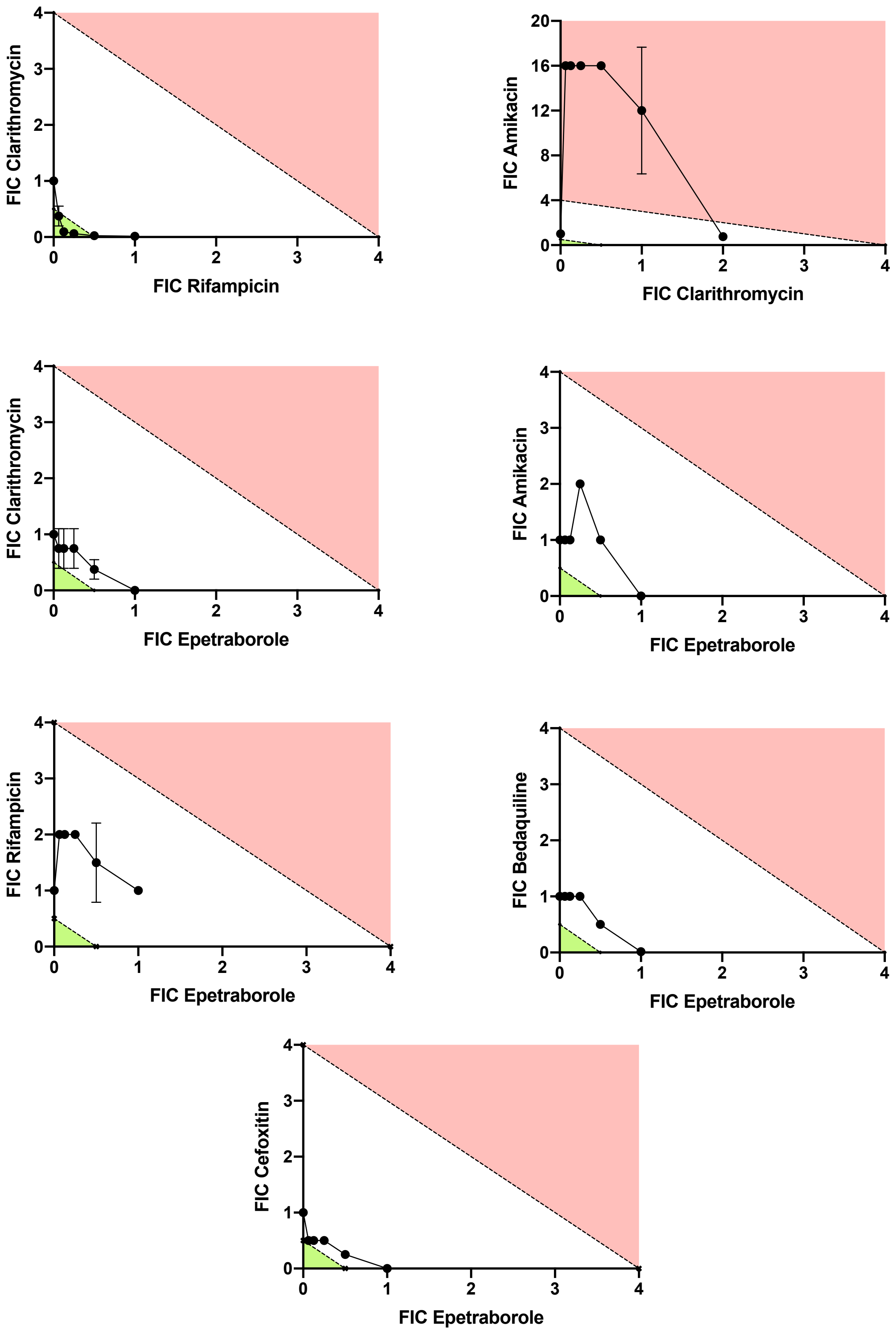


**Supplementary Figure 4.**
Isobolograms for potential combination therapy with EPT. Green area indicates synergy (FICI < 0.5); red area indicates antagonism (FICI ≥ 4.0); white area indicates indifferent. RIF and CLR, and AMK and CLR are used as synergy and antagonism controls, respectively. Data shown is from checkerboard assays done in technical duplicate.

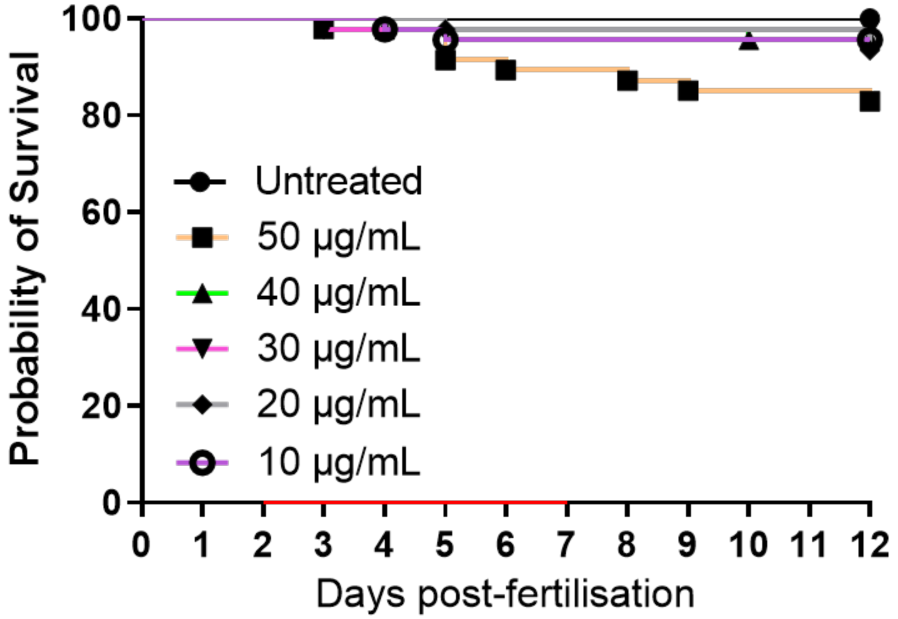


**Supplementary Figure 5.**
EPT toxicity against zebrafish embryos. Groups of uninfected embryos were immersed in water containing 10 to 50$\mu$g/mL EPT for 5 days. Red bar indicates duration of treatment. Data is from two independent experiments.

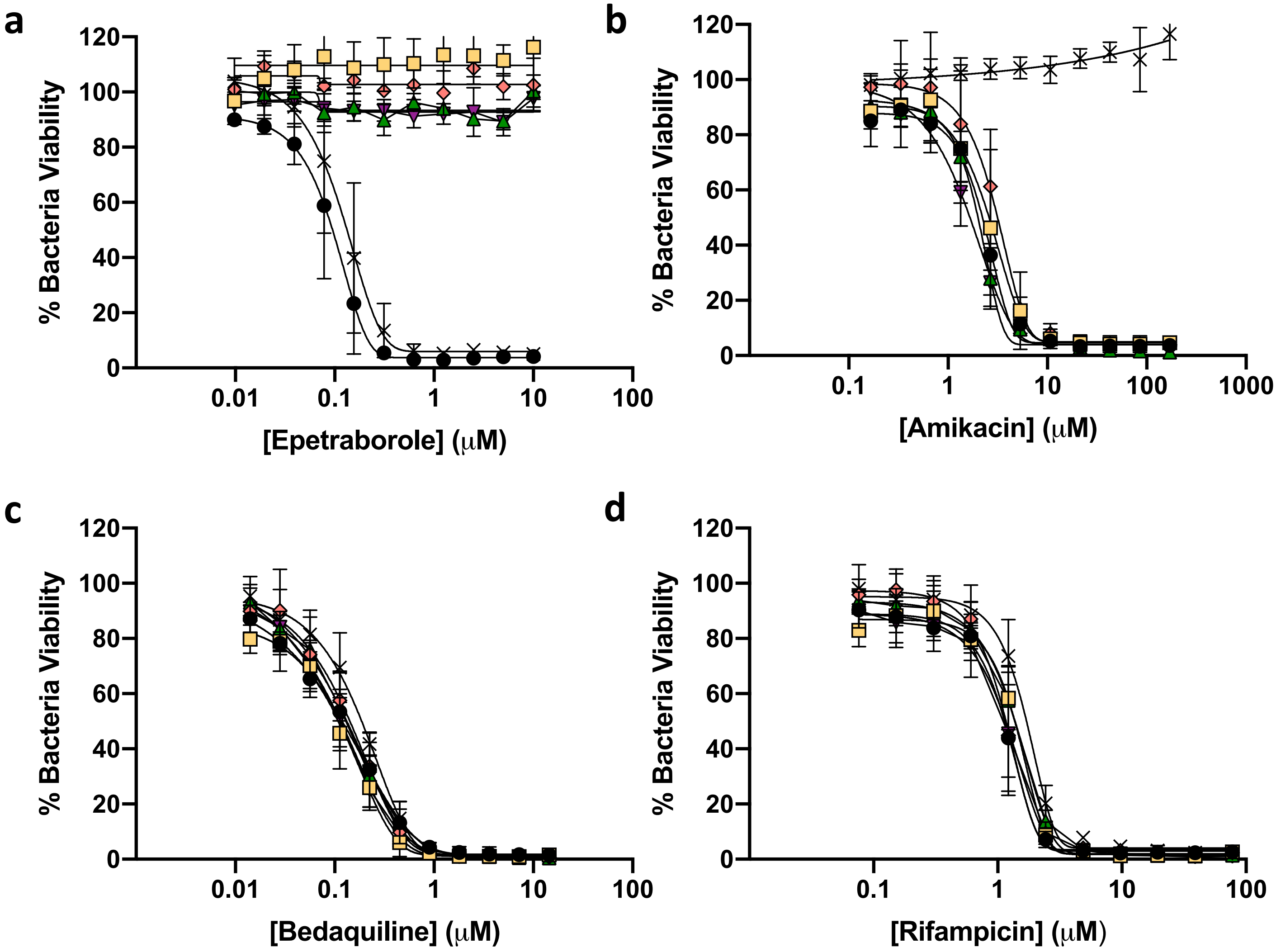


**Supplementary Figure 6.**
Drug susceptibility assessment of resistant mutants. **a-d** Dose-response curves of isolated EPT (10X, 20X, 40X MIC) mutants or AMK (40X MIC) mutant. AMK was used as control compound for mutant isolation. RIF and BDQ were used for cross-resistance verification. ATCC 19977 (black circles), D436H mutant-1 (yellow squares), D436H mutant-2 (pink diamonds), D436H mutant-3 (green triangles), D436H mutant-4 (purple inverted triangles), AMK 40X (crosses). Data is mean ± SD from technical triplicates.

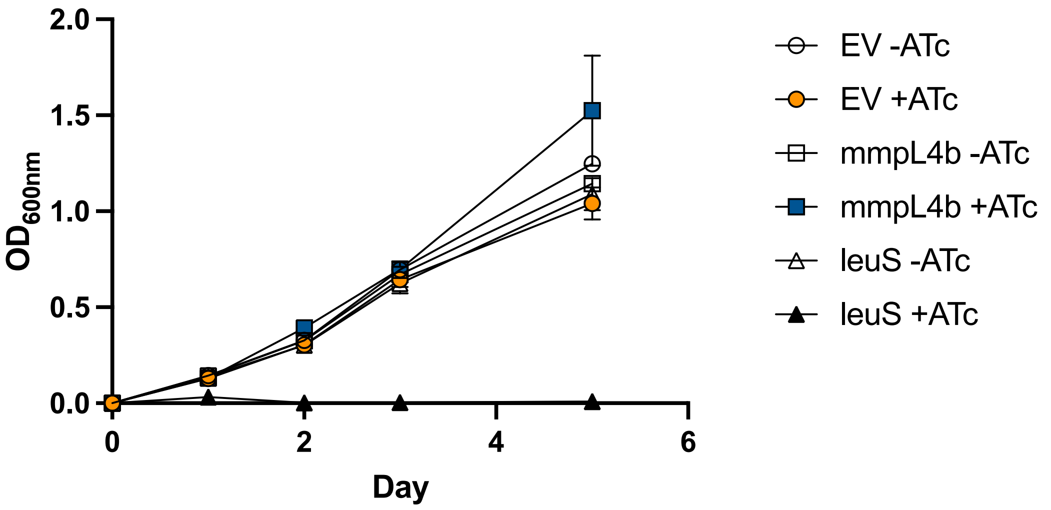


**Supplementary Figure 7.**
*leuS* is an essential gene in *M. abscessus* using CRISPRi gene knockdown. *M. abscessus* carrying integrated empty CRISPRi vector (EV, circles), CRISPRi:*mmpL4b* as a non-essential gene control (squares), or CRISPRi:*leuS* (triangles) were grown with/without 2 μM ATc. Data is mean ± SD from biological triplicates.

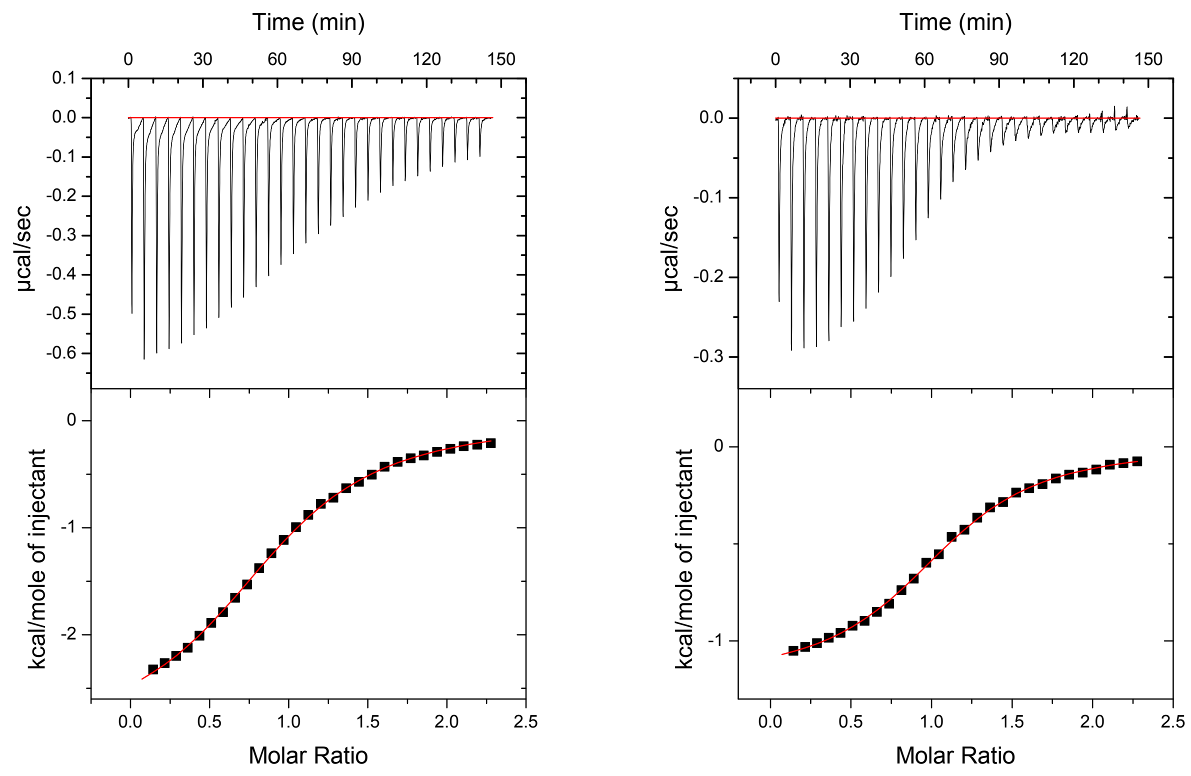


**Supplementary Figure 8.**
Thermodynamic analysis of EPT binding to LeuRS editing domain. Heat of injection (upper panel) and single-site binding model of the integrated isotherm (lower panel). *Mabs* LeuRS (left) or *Mtb* LeuRS (right) editing domains bound to EPT.

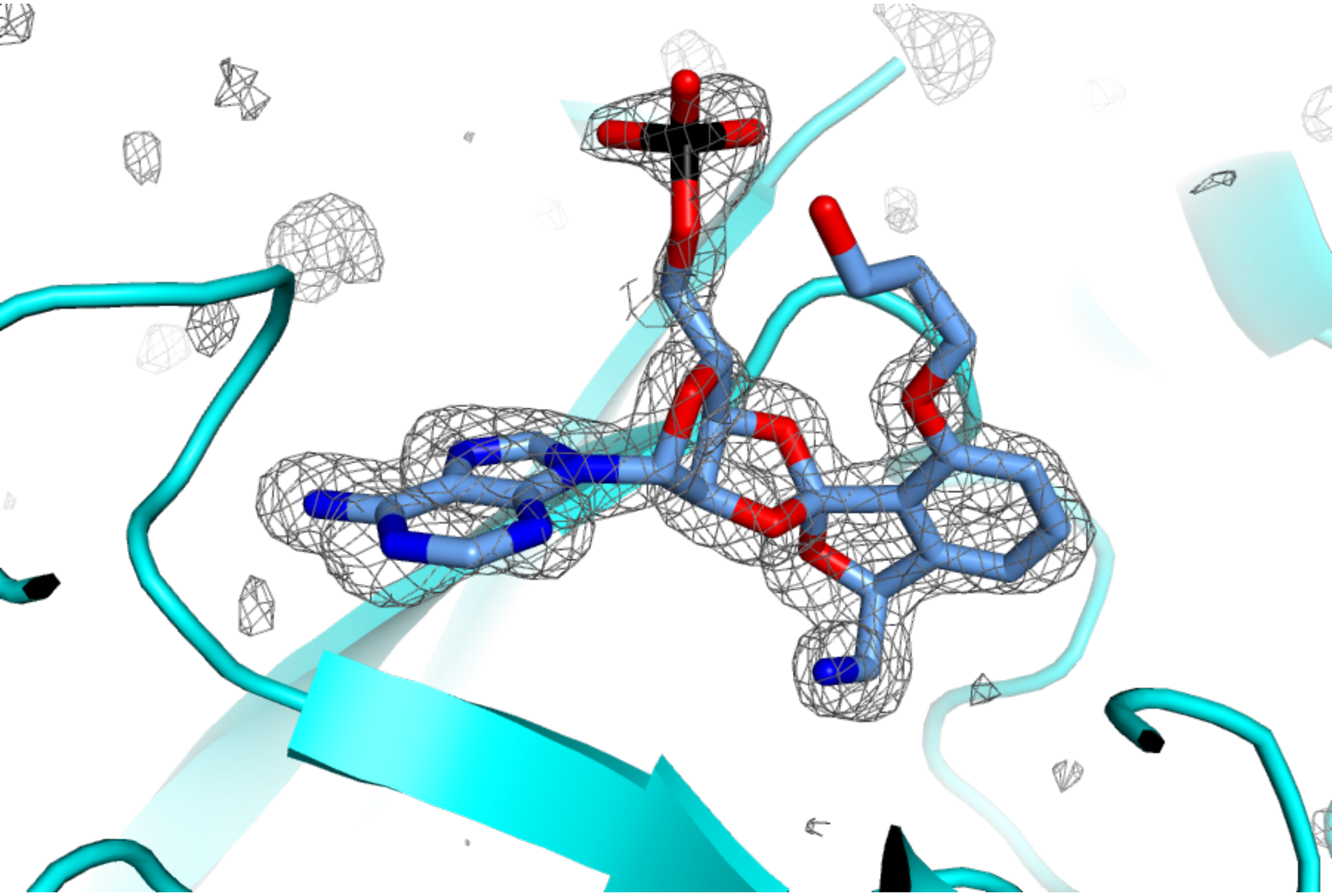

**Supplementary Figure 9.**
Difference maps for the EPT-AMP adduct in 7N12. Unbiased FO-FC difference maps, calculated with phases from a model that never included ligand.

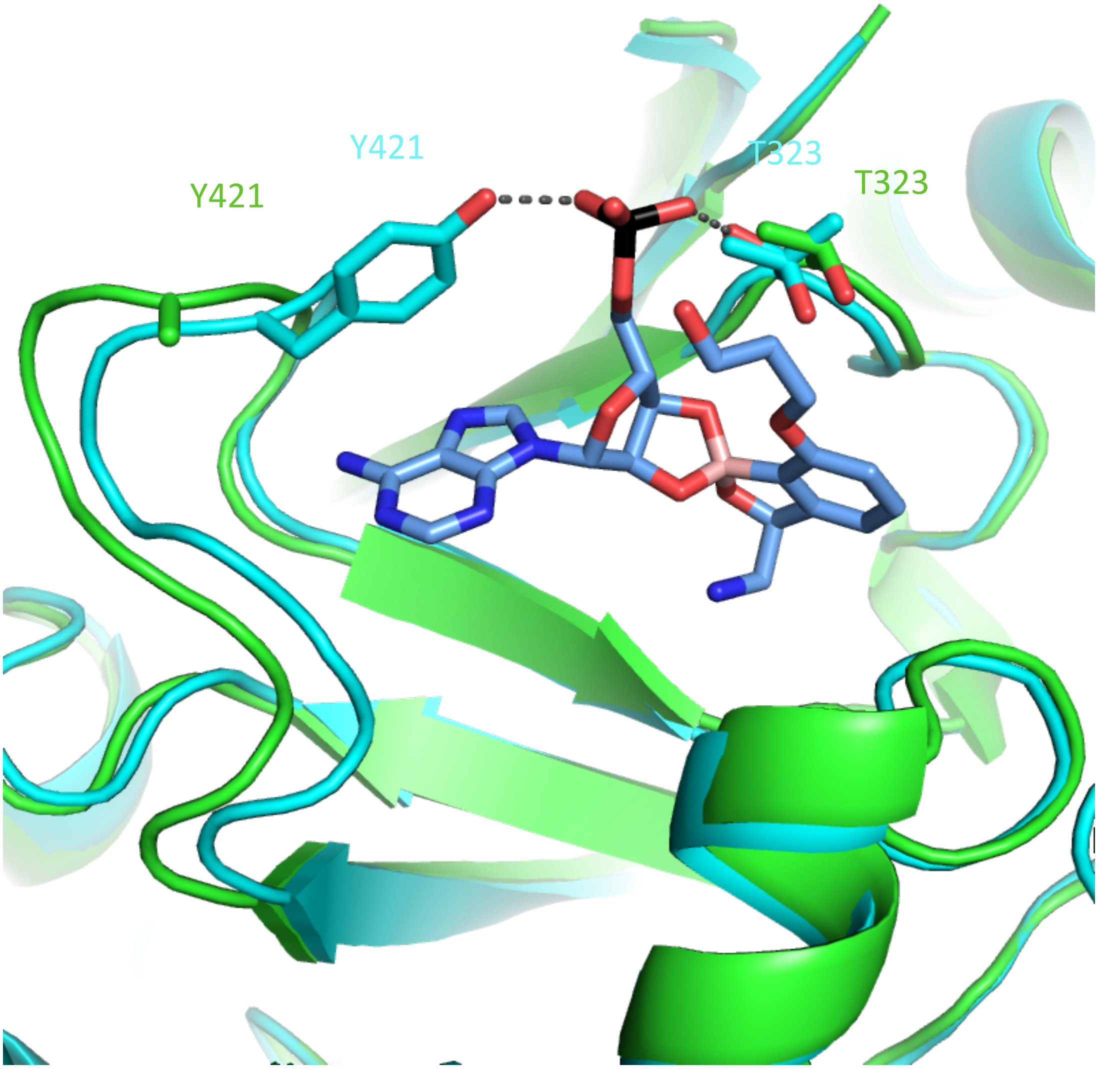


**Supplementary Figure 10.**
Comparison of apo (green, PDB 7N11) and co-complex (cyan, PDB 7N12) structures of *M. abscessus* LeuRS bound to EPT-AMP.

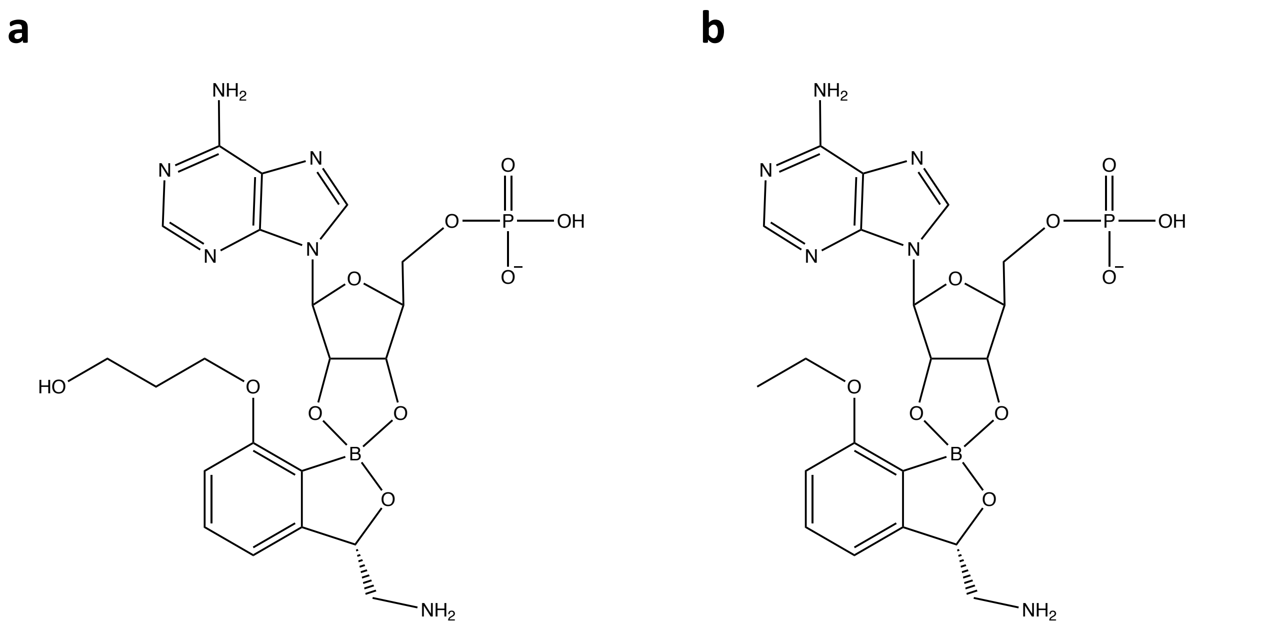


**Supplementary Figure 11.**
Benzoxaborole inhibitors of LeuRS. **a** EPT-AMP adduct bound to *M. abscessus* LeuRS. **b** BNZ-AMP adduct bound to *M. tuberculosis* LeuRS.

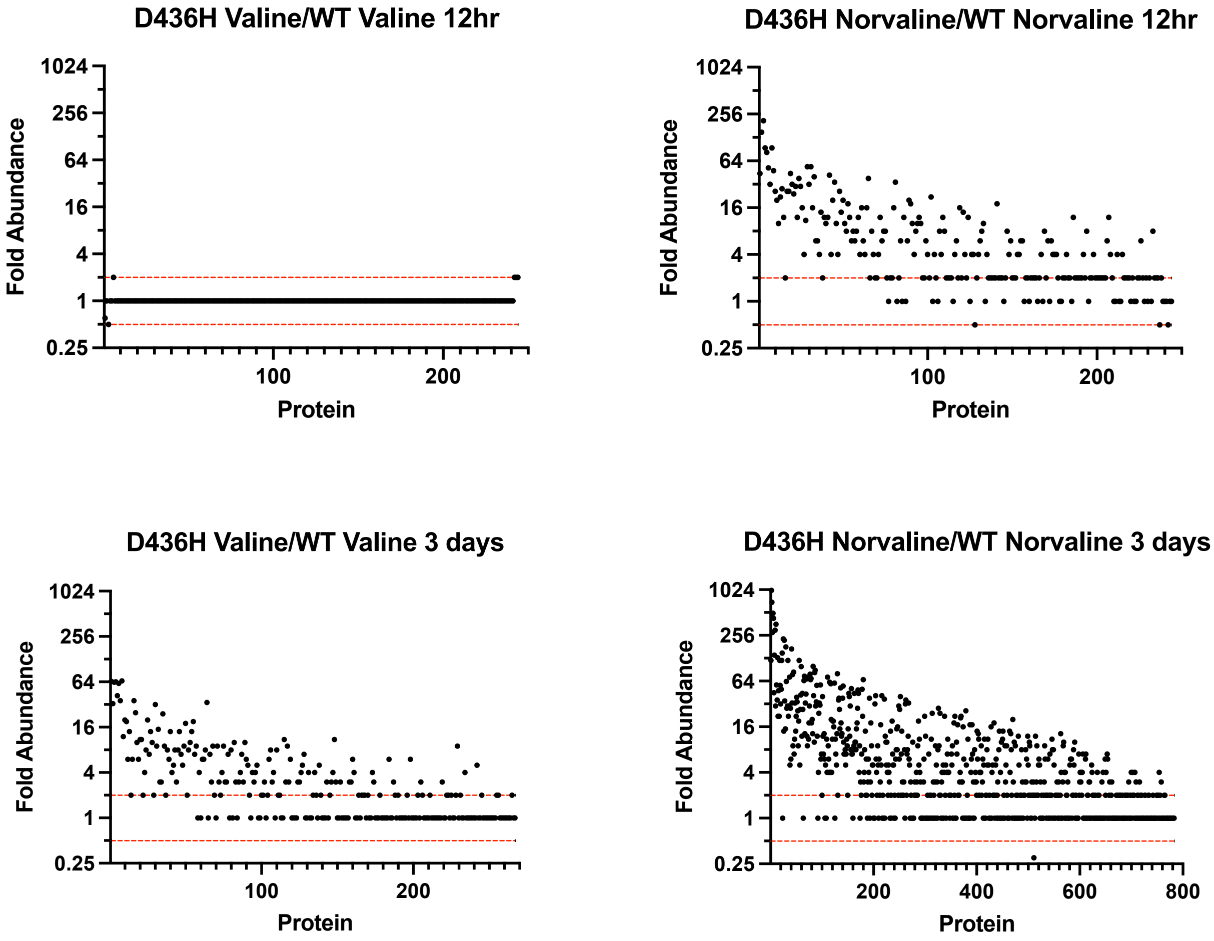


**Supplementary Figure 12.**
Norvaline stress alters the proteome in editing deficient mutants. ATCC 19977 or D436H mutant were grown in 0.5mM norvaline or 0.5mM valine for 12 hours or 3 days. Valine was used as a specificity control. Total cell lysate was collected. Protein fold abundance shown as D436H mutant relative to wild-type *M. abscessus*. Dashed red lines indicate 0.5x and 2x fold abundance thresholds for biologically relevant changes.

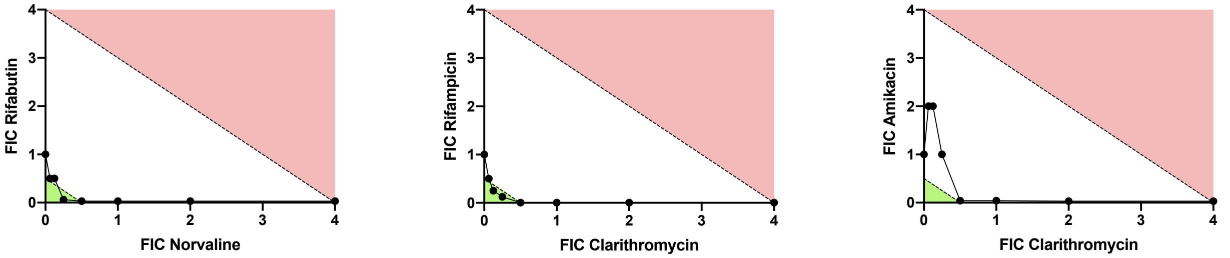


**Supplementary Figure 13.**
Norvalination of the proteome leads to synergy with RFB. Isobolograms for potential combination therapy with norvaline. Green area indicates synergy (FICI < 0.5); red area indicates antagonism (FICI ≥ 4.0); white area indicates indifferent. Data shown is mean ± SD from three independent biological replicates.

**Supplementary Table 1. MMV Open pandemic response box hits and reference compounds**

| Parameter | Compound | | | | |
|---|---|---|---|---|---|
| | EPT | ERV | IQN | BDQ | AMK |
| $MIC_{90}$ (µM) | 0.30 | 3.0 | 8.4 | 0.90 | 6.8 |
| $TD_{50}$ (µM) | >100 | >100 | >100 | 17[a] | 30 |
| TI ($TD_{50}$/$MIC_{90}$) | >330 | 33 | 12 | 19 | 4.4 |
| MW (g/mol) | 237.06 | 631.52 | 393.40 | 555.51 | 585.60 |

EPT, Epetraborole; ERV, Eravacycline; IQN, Isoquinoline urea; BDQ, Bedaquiline; AMK, Amikacin; $MIC_{90,}$ concentration of drug that inhibits 90% of bacterial growth; $TD_{50}$, concentration of drug with 50% toxicity. TI, therapeutic index.

[a]Data from Lupien, A et al. *Antimicrob Agents Chemother*. **2018**

**Supplementary Table 2. Carbon dependent variability on EPT activity**

| | Media | | | |
|---|---|---|---|---|
| | 7H9 G + Tw80 | 7H9 G - Tw80 | 7H9 A + Tw80 | CaMH + Tw80 |
| $MIC_{90}$ (µM) | 0.38 ± 0.06 | 0.04 ± 0.03 | 0.06 ± 0.02 | 2.0 ± 0.8 |

G, Glycerol; A, Acetate; Tw80, Tween-80; CaMHB, Cation-adjusted Muller-Hinton

**Supplementary Table 3. Epetraborole *in vitro* activity against clinical isolates**

| | | | MIC (µM) | | | | | |
|---|---|---|---|---|---|---|---|---|
| **Isolate** | **Subspecies** | **Morphology[a]** | **EPT** | **AMK** | **RIF** | **BDQ** | **CFX** | **CLR** |
| ATCC 19977 | *abscessus* | S | 0.23 | 8.2 | 11 | 1.2 | 28 | 1.2 |
| ATCC 19977 | *abscessus* | R | 0.40 | 4.1 | 2.2 | 0.49 | 28 | 0.87 |
| MT16_1065 | *abscessus* | S | 0.15 | 3.6 | >29 | 0.65 | 31 | 4.5 |
| MT16_6490 | *abscessus* | S | 0.11 | 9.7 | >29 | 1.5 | 30 | 3.9 |
| MT15_1748 | *massiliense* | R | 0.080 | 24 | 11 | 0.41 | 61 | 0.091 |
| MT15_1749 | *massiliense* | R | 0.051 | 15 | 8.1 | 0.72 | 37 | 0.11 |
| Paris 167 | *bolletii* | R | 0.099 | 8.2 | 9.0 | 0.70 | 37 | 3.7 |
| AV | *bolletii* | S | 0.17 | 12 | >29 | 2.3 | 35 | 0.20 |

[a]Morphology as determined by smooth (S) or rough (R) colonies on 7H10 agar. Drugs used: EPT, Epetraborole; AMK, Amikacin; RIF, Rifampicin; BDQ, Bedaquiline; CFX, Cefoxitin; CLR, Clarithromycin

**Supplementary Table 4. Epetraborole *in vitro* activity spectrum**

| | | MIC (µM) | | | | | |
|---|---|---|---|---|---|---|---|
| **Species** | **Gram[a]** | **EPT** | **AMK** | **RIF** | **BDQ** | **CFX** | **CLR** |
| *M. abscessus* | AF | 0.23 | 8.2 | 11 | 1.2 | 28 | 1.2 |
| *M. avium hominissuis* | AF | 9.9 | 9.6 | 0.077 | 0.068 | 11 | 0.24 |
| *M. avium intracellulaire* | AF | 9.9 | 6.5 | 0.077 | 0.025 | 0.98 | 0.019 |
| *M. tuberculosis H37Rv* | AF | 1.7 | 0.51 | 0.0077 | 0.47 | >230 | >1.7 |
| *M. tuberculosis Erdman* | AF | 0.69 | 0.82 | 0.0079 | 0.11 | >230 | >1.7 |
| *B. cereus* | + | >9.9 | 110 | <0.077 | >14 | >230 | >11 |
| *C. glutamicum* | + | 7.3 | 0.11 | <0.077 | >14 | 230 | >11 |
| *E. coli* | - | 2.1 | 0.26 | 4.7 | >14 | 9.4 | >11 |
| *P. aeruginosa* | - | >9.9 | 0.38 | 18 | >14 | >230 | >11 |

[a]Acid fast (AF); gram positive (+); gram negative (-). EPT, Epetraborole; AMK, Amikacin; RIF, Rifampicin; ERV, BDQ, Bedaquiline; CFX, Cefoxitin; CLR, Clarithromycin. Species: *Mycobacterium abscessus*, *Mycobacterium avium hominissuis*, *Mycobacterium avium intracellulaire*, *Mycobacterium tuberculosis* H37Rv, *Mycobacterium tuberculosis* Erdman, *Bacillus cereus*, *Corynebacterium glutamicum*, *Escherichia coli*, *Pseudomonas aeruginosa*

**Supplementary Table 5. *In vitro* resistance frequency**

| | Resistance frequency[a] | | |
|---|---|---|---|
| **Compound[b]** | **10X** | **20X** | **40X** |
| Amikacin | $1.3 \times 10^{-8}$ | N.D | N.D |
| Epetraborole | $2 \times 10^{-9}$ | $1 \times 10^{-9}$ | $2 \times 10^{-9}$ |

[a]N.D: not determined. [b]$MIC_{90}$ values on 7H10 agar used in this experiment were 4µg/mL and 0.25µgmL for amikacin and epetraborole, respectively.

**Supplementary Table 6. Primers**

| Primers | Function | Sequence | Source |
|---|---|---|---|
| 1. pMV306hsp60_leuS_F | Cloning | GATGATCTGCAGAACCGAAACCCAGCACGACG | This study |
| 2. pMV306hsp60_leuS_R | Cloning | GATGATAAGCTTCTAGACGACCAGGTTCACCATG | This study |
| 3. pMV306hsp60_leuS_F1 | Sequencing | GTAAGTAGCGGGGTTGCCGT | This study |
| 4. pMV306hsp60_leuS_R1 | Sequencing | GCTGCCGCTTGTAGTTGACG | This study |
| 5. pMV306hsp60_leuS_F2 | Sequencing | TGCAGACCGGCACCCATCC | This study |
| 6. pMV306hsp60_leuS_R2 | Sequencing | TTTGACCTTGTCCGGCCAGT | This study |
| 7. pMV306hsp60_leuS_F3 | Sequencing | CGCCTACTCCGACAGGTTGA | This study |
| 8. pMV306hsp60_leuS_R3 | Sequencing | CCGGCAAACCGAAGGTGTT | This study |
| 9. pMV306hsp60_leuS_F4 | Sequencing | ATGGCACCGGTGCCATCAT | This study |
| 10. pMV306hsp60_leuS_R4 | Sequencing | GCGGCATCACGTTGGTGTC | This study |
| 11. pMV306hsp60_leuS_F5 | Sequencing | AATGTCGAGCTGGACCTCGG | This study |
| 12. pMV306hsp60_leuS_R6 | Sequencing | GCAGCGTATCGGCACCATAGT | This study |
| 13. pMV306hsp60_leuS_F6 | Sequencing | GCCTCAAGAACTCGATCTCGC | This study |
| 14. pMV306hsp60_leuS_R6 | Sequencing | TGGCAGTCGATCGTACGCTAG | This study |
| 15. CRISPRi_leuS_sgRNA_F | CRISPRi | GGGAACTCTTGTTGCACCTCTGCGCCCTG | This study |
| 16. CRISPRi_leuS_sgRNA_R | CRISPRi | AAACCAGGGCGCAGAGGTGCAACAAGAGT | This study |

**Supplementary Table 7. Thermodynamic analysis of EPT binding with LeuRS**

| Bacteria | ΔG (kcal mol$^{-1}$) | ΔH (kcal mol$^{-1}$) | -TΔS (kcal mol$^{-1}$) | Kd ($\mu$M) |
|---|---|---|---|---|
| *M. abscessus* | -6.49 ± 0.07 | -3.2 ± 0.4 | -3.3 ± 0.3 | 16 ± 4 |
| *M. tuberculosis* | -6.9 ± 0.2 | -1.1 ± 0.3 | -5.5 ± 0.5 | 10 ± 4 |

ΔG, change in Gibbs free energy; ΔH, change in enthalpy; ΔS, change in entropy; T, temperature (303K) Kd, dissociation constant

**Supplementary Table 8. Data collection and refinement statistics**

| | *M. abscessus* LeuRS editing domain | *M. abscessus* LeuRS editing domain in complex with epetraborole-AMP adduct |
|---|---|---|
| **Data collection** | | |
| Space group | $P\ 2_1\ 2_1\ 2_1$ | C 1 2 1 |
| Cell dimensions | | |
| *a*, *b*, *c* (Å) | 37.25, 51.59 116.15 | 113.439, 37.1074, 100.287 |
| α, β, γ (°) | 90, 90, 90 | 90, 112.279, 90 |
| Resolution (Å) | 1.69-50.00 (1.69-1.72) * | 1.52-92.97 (1.52-1.55) * |
| $R_{sym}$ | 0.263 (0.824) * | 0.100 (1.071) * |
| $I / \sigma I$ | 4.4 (0.5) * | 7.7 (0.3) * |
| Completeness (%) | 76.8 (5.5) * | 94.21 (49.23) * |
| Redundancy | 6.8 (1.1) * | 5.54 (3.08) * |
| **Refinement** | | |
| Resolution (Å) | 2.1 | 1.7 |
| No. reflections | 14741 | 109484 |
| $R_{work} / R_{free}$ | 0.1841 / 0.2258 | 0.1843 / 0.2265 |
| No. atoms | 2968 | 5964 |
| Protein | 2822 | 5435 |
| Ligand | / | 126 |
| Ion | 15 | 5 |
| Water | 131 | 384 |
| *B*-factors | | |
| Protein | 25.44 | 20.14 |
| Ligand | / | 19.98 |
| Ion | 63.20 | 56.13 |
| Water | 31.17 | 31.70 |
| R.m.s. deviations | | |
| Bond lengths (Å) | 0.008 | 0.013 |
| Bond angles (°) | 0.999 | 1.216 |
| **PDB accession code** | 7N11 | 7N12 |
